# Local postsynaptic signalling on slow time scales in reciprocal olfactory bulb granule cell spines matches asynchronous release

**DOI:** 10.1101/2020.09.03.281642

**Authors:** Tiffany Ona Jodar, Vanessa Lage-Rupprecht, Nixon M. Abraham, Christine R. Rose, Veronica Egger

## Abstract

In the vertebrate olfactory bulb (OB), axonless granule cells (GC) mediate self- and lateral inhibitory interactions between mitral/tufted cells via reciprocal dendrodendritic synapses. Locally triggered release of GABA from the large reciprocal GC spines occurs on both fast and slow time scales, possibly enabling parallel processing during olfactory perception. Here we investigate local mechanisms for asynchronous spine output.

To reveal the temporal and spatial characteristics of postsynaptic ion transients, we imaged spine and adjacent dendrite Ca^2+^- and Na^+^-signals with minimal exogenous buffering by the respective fluorescent indicator dyes upon two-photon uncaging of DNI-glutamate in OB slices from juvenile rats. Both postsynaptic fluorescence signals decayed slowly, with average half durations in the spine head of t_1/2__Δ[Ca^2+^]_i_ ~500 ms and t_1/2__Δ[Na^+^]_i_ ~1000 ms. We also analysed the kinetics of already existing data of postsynaptic spine Ca^2+^-signals in response to glomerular stimulation in OB slices from adult mice, either WT or animals with partial GC glutamate receptor deletions (NMDAR: GluN1 subunit; AMPAR: GluA2 subunit). In a large subset of spines the fluorescence signal had a protracted rise time (average time to peak ~400 ms, range 20 ms - >1000 ms). This slow rise was independent of Ca^2+^ entry via NMDARs, since similarly slow signals occurred in ΔGluN1 GCs. Additional Ca^2+^ entry in ΔGluA2 GCs (with AMPARs rendered Ca^2+^-permeable), however, resulted in larger ΔF/Fs that rose yet more slowly.

Thus GC spines appear to dispose of several local mechanisms to promote asynchronous GABA release, which are reflected in the time course of mitral/tufted cell recurrent inhibition.

## Introduction

In the vertebrate olfactory bulb (OB), the lateral dendrites of the principal mitral and tufted cells are interconnected via local GABAergic interneurons. The most abundant class of these local neurons, the axonless granule cells (GC), mediate self- and lateral inhibitory interactions between mitral/tufted cells via reciprocal dendrodendritic synapses that on the GC dendrite are housed in large spines. These reciprocal synapses have been directly demonstrated to play a role in odor discrimination and learning (Abraham et al., 2010; Gschwend et al., 2015). Moreover, they are also critically involved in generating bulbar γ-oscillations (Rall and Shepherd, 1968; Nusser et al., 2001; Lagier et al., 2004), which are thought to contribute to odor coding via synchronization and gating of mitral cell output (e.g. Buonviso et al., 2003; Fukunaga et al., 2014; Osinsky and Kay, 2016).

Recordings of dendrodendritic inhibition (DDI) of mitral cells have revealed that recurrent inhibition happens as a barrage of IPSCs (Schoppa et al., 1998; Isaacson and Strowbridge, 1998). Within this barrage, early IPSCs will occur with a very short latency (below 10 ms), but recurrent activity takes several hundreds of milliseconds to subside. While this long tail of recurrent inhibition is unlikely to directly contribute to odor discrimination itself (Abraham et al., 2004, 2010; Uchida and Mainen, 2003), it may well play a role in learning and memory formation (Gschwend et al., 2015; see Discussion).

The underlying asynchronous release is at least to a major extent due to processing in GCs, since asynchronous responses were demonstrated following flash photolysis of Ca^2+^ in mitral cell lateral dendrites (Chen et al., 2000). Moreover, while the massive release of glutamate during the commonly used protocol for mitral cell excitation (20-50 ms depolarization in the voltage-clamp mode) might result in activation of slow release pathways not accessible to unitary transmission, we have shown recently, that local, unitary-like two-photon uncaging of glutamate (TPU) can still cause prolonged release of GABA within a time window of up to 500 ms post uncaging (Lage-Rupprecht et al., 2020).

As to possible mechanisms for late output, unitary EPSPs evoked by spontaneous mitral/tufted cell input or local TPU are mediated by both AMPA and NMDA receptors and decaying with a time constant < 50 ms (as recorded at the GC soma, Bywalez et al., 2015). Thus slower actions downstream of ionotropic receptors would be required to trigger cascades that result in asynchronous release events beyond 100 ms. A number of **global** mechanisms has been proposed to promote asynchronous release from GC spines. These include a delay of global GC action potentials (AP) due to the prominent I_A_ current (Schoppa and Westbrook, 1999; Kapoor and Urban, 2006), and a prolonged Ca^2+^ entry due to synaptic activation of a non-specific cation current I_CAN_, possibly in coincidence with global APs (Hall and Delaney, 2002, Egger, 2008; Stroh et al., 2012).

Since as in other synapses reciprocal release of GABA is Ca^2+^-dependent (Isaacson and Strowbridge, 1998), how could spine Ca^2+^ signals mediate asynchronous release? To answer this question, we explored several potential **local** mechanisms that might be involved in slow spine Ca^2+^ signalling and thus are directly related to the biophysical properties of individual spines.

While the endogenous Ca^2+^ buffering capacity κ_E_ in GC spines is not unusually high (~120) and thus cannot explain lingering Ca^2+^, the Ca^2+^ extrusion from the spine cytosol is sluggish (rate γ ~500 s^−1^ at RT), which might support asynchronous output (Egger and Stroh, 2009). As to postsynaptic spine Ca^2+^ signals upon glomerular mitral cell stimulation (100 μM OGB-1, Egger et al., 2005), responses in juvenile rat GC spines are robust, with an average amplitude of ~40 % ΔF/F, and rise within ~80 ms. Their decay kinetics are slower than those of backpropagating AP-mediated transients, with a clearly bimodal distribution of durations. While ~2/3 of spine signals decayed by half within ~600 ms, the remaining 1/3 decayed very slowly, with half durations beyond 1.5 s. Identical signal properties including the fraction of ‘slow spines’ are observed in response to TPU of glutamate (Bywalez et al., 2015). This unitary postsynaptic Ca^2+^ entry is mainly mediated by NMDA receptors, with additional contributions by low- and high voltage activated Ca^2+^ channels and Ca^2+^-induced Ca^2+^ release (CICR; Egger et al., 2005). Relief from the Mg^2+^ block is provided via a local AP (‘spine spike’, Bywalez et al., 2015), while native GC AMPA receptors are not Ca^2+^-permeable (Jardemark et al., 1992; Egger et al., 2005).

However, these previous recordings of synaptic fluorescence transients were all performed with the Ca^2+^ indicator dye OGB-1 (100 μM, K_d_ = 200 nM). Therefore transients are substantially buffered and do not reflect true kinetics of Ca^2+^ signals (Egger and Stroh, 2009), even though their observed decay times seem to fit well with the time scale of asynchronous release. Thus here we asked whether synaptic signals are indeed slow, also in comparison to AP-mediated transients, by using a low-affinity Ca^2+^ dye.

Moreover, we had observed earlier on that the triggering of long-lasting depolarizations in the wake of synaptically evoked GC APs required both NMDA receptor activation and opening of voltage-gated Na^+^ channels (Na_v_; observed in both juvenile rats and adult mice) and that these long-lasting depolarizations were carried by TRPC1/4 heteromeric channels (Egger, 2008; Stroh et al., 2012). Since we know by now that local postsynaptic inputs can trigger ‘spine spikes’ within the spine head (Bywalez et al., 2015), we hypothesized that already local synaptic signals might involve a long-lasting cationic inward current I_CAN_ (via TRPCs or otherwise). To this end, we combined Na^+^ imaging based on the dye SBFI (Rose et al., 1999; Ona-Jodar et al., 2017) with TPU of glutamate.

Finally, we noticed that in spite of similar endogenous Ca^2+^ dynamics and similar amplitudes of spontaneous and evoked synaptic transients in adult mouse and juvenile rat GCs (Egger et al., 2005; unpublished observations Egger and Stroh, Egger and Stroh, 2009; Abraham et al., 2010), postsynaptic Ca^2+^ signals in adult mouse GCs were yet slower with regard to their rise time (original data set from Abraham et al. 2010, in which kinetics had not been analysed quantitatively). This study had revealed a correlation between behavioural performance in a go/no-go odor discrimination task, and modifications of postsynaptic ΔCa^2+^ into the majority of GC spines via viral transfection, across three sample groups: (1) WT mice, (2) mice with a deletion of the GluA2 AMPAR subunit (ΔGluA2; increased Ca^2+^ entry) which resulted in faster discrimination and thus a gain of function, and (3) mice with a deletion of the NR1 NMDAR subunit (ΔGluN1, i.e. reduced Ca^2+^ entry), which resulted in slowed discrimination, i.e. a loss of function. Here we provide a quantitative analysis of the kinetics of the respective Ca^2+^ signals and the response probability and use the genetic pharmacology provided by the viral knockdown to infer possible mechanisms for the slow rise.

In summary, here we aim to unravel the pyhsiological time courses of postsynaptic Ca^2+^ and Na^+^ signals in juvenile rat GCs in order to investigate their overlap with the previously established time course of asynchronous release. Moreover, we describe an additional potential source of delayed release from adult mouse GC spines.

## Methods

### Juvenile rat experiments: Preparation, electrophysiology

Sagittal olfactory bulb brain slices (thickness 300 μm) were prepared in artificial cerebrospinal fluid (ACSF, composition see below) following procedures in accordance with the rules laid down by the EC Council Directive (86/89/ECC) and German animal welfare legislation. Slices were incubated a water bath at 33°C for 30 min and then kept at room temperature (22°C) until recordings were performed.

The extracellular ACSF was bubbled with carbogen and contained (in mM): 125 NaCl, 26 NaHCO3, 1.25 NaH2PO4, 20 glucose, 2.5 KCl, 1 MgCl2, and 2 CaCl2. Whole cell current clamp recordings were performed at room temperature (22°C) and granule cells were held near their resting potential of −80 mV. Granule cells were filled with an internal solution containing the following substances (in mM): 130 K-Methylsulfate, 10 HEPES, 4 MgCl_2_, 2 ascorbic acid, 10 phosphocreatine-di-tris-salt, 2.5 Na_2_ATP, 0.4 NaGTP, and 1 mM SBFI (Na^+^-binding benzofuran isophthalate, Teflabs, Austin, TX and Molecular Probes, Eugene, OR, USA) or 0.1 OGB-6F (Ca^2+^ indicator, Thermofisher Scientific, Waltham. Massachusetts, USA). The patch pipette resistance varied between 6 and 7 MΩ.

### Juvenile rat experiments: Combined two-photon imaging and uncaging

For Na^+^ imaging experiments, electrophysiology and imaging were performed as in (Ona-Jodar et al., 2017), and for Ca^2+^ imaging experiments as in (Bywalez et al., 2015). Uncaging is also described in detail in (Bywalez et al., 2015). Imaging and uncaging were performed on a Femto-2D-uncage microscope (Femtonics, Budapest, Hungary). Two tunable, Verdi-pumped Ti:Sa lasers (Chameleon Ultra I and II respectively, Coherent, Santa Clara, CA, USA) were used in parallel. The first laser was set either to 840 nm for excitation of OGB-6F or to 800 nm for excitation of SBFI in GC spines and dendrites, and the second laser was set to 750 nm for uncaging of caged glutamate. The two laser lines were directly coupled into the pathway of the microscope with a polarization cube (PBS102, Thorlabs Inc, Newton, NJ, USA) and two motorized mirrors. As caged compound we used DNI-caged glutamate (DNI; Femtonics). DNI was used at 1 mM in a closed perfusion circuit with a total volume of 12 ml. Caged compounds were washed in for at least 10 minutes before starting measurements. The uncaging laser was switched using an electro-optical modulator (Pockels cell model 350-80, Conoptics, Danbury, CT, USA).

Na^+^ and Ca^2+^ signals were imaged in line scanning mode with a temporal resolution of ~1 ms. The scan position was checked and readjusted if necessary before each measurement to account for drift.

### Adult mouse GC Ca^2+^ imaging (data from Abraham et al. 2010)

The experiments in adult mice are described in Abraham et al. 2010; the new analyses presented here are based on the very same data set. Briefly (see Abraham et al. 2010 for details), GC-specific deletion of GluA2 AMPAR subunit and GluN1 NMDAR subunit had been achieved by viral expression of Cre recombinase in mice with conditional alleles of GluA2 and GluN1. To restrict the deletion to GCs we had injected rAAV Cre only in the anterior portion with respect to the center of the dorsal OB surface. OB slices had been prepared after an incubation time of at least 2 weeks. GluA2- and GluN1-depleted GCs had been identified by somatic fluorescence arising from co-expression of Cre recombinase and Kusabira orange (Tang et al., 2009). Mitral/tufted cells had been activated via glomerular extracellular stimulation, and responding GC spines had been searched for with two-photon Ca^2+^ imaging in GCs patched below the stimulated glomerulus that had responded to glomerular stimulation with a detectable EPSP (see also Fig. 3A).

### Data analysis and statistics

Imaging data were analysed with custom written macros in Igor Pro (Wavemetrics, Lake Oswego, OR, USA), as described previously (Egger et al., 2003, 2005). All imaging signals (OGB-6, OGB-1, SBFI) were analysed in terms of ΔF/F = (F(t)-F_0_)/F_0_. Rise times were measured between 20% and 80% of the absolute maximal ΔF/F amplitude, and half durations t_1/2_ reflect the period from this maximal amplitude to the half-maximal amplitude. SBFI ΔF/F signals were converted into absolute concentration changes Δ[Na^+^]_i_ according to the previously established calibration on the same system: for non-saturating signals a 10% change in fluorescence emission of SBFI corresponds to a change of 22.3 mM in [Na^+^]_i_ (Ona-Jodar et al., 2017). The response probability is an estimate of the release probability and was calculated as the ratio of detected responses to the total number N of stimulations (average N = 14 ± 5 in WT).

Statistical comparisons were made with non-parametric tests (Wilcoxon test for paired and Mann-Whitney test for unpaired data sets). Comparisons between WT GC responses and the ΔGluA2 and ΔGluN1 GC groups were made via pairwise Mann-Whitney tests with Bonferroni correction. Frequency distributions of parameters were compared with the Kolmogorov-Smirnov test. Mean values are given ± S.D.

## Results

### Time course of synaptic spine Ca^2+^ signals with minimal exogenous buffering

To investigate the local mechanisms underlying the asynchronous component of reciprocal GABA release, we aimed to detect local postsynaptic Ca^2+^ signalling in GC spines with as little exogenous buffering as possible, since sluggish extrusion of Ca^2+^ might also contribute to delayed release (Egger and Stroh, 2009). The low-affinity dye OGB-6F (K_d_ ≈ 8 μM, Tran et al., 2018) was used at a concentration of 100 μM, where the kinetics of OGB-6F fluorescence transients in response to single somatic APs (ΔF/F)_sAP_ are identical to the kinetics determined by extrapolation of measurements with varying concentrations of OGB-1 (K_d_ ≈ 0.2 μM) to zero added buffer (Egger and Stroh, 2009).

TPU of DNI with similar parameters as in Bywalez et al. (2015; Fig. 1A) evoked Ca^2+^ transients (ΔF/F)_TPU_ in juvenile rat GC spine heads (postnatal days PND11-19), with a mean amplitude of 24 ± 9 %, a mean rise time of 55 ± 32 ms and a mean half duration t_1/2_ of 445 ± 225 ms (n = 11 spines, Fig. 1B, C). These transients were strictly localized to the spine head (mean (ΔF/F)_TPU_ amplitude in adjacent dendritic shaft 2 ± 1 %, ratio vs spine head 0.09 ± 0.03). While t_1/2_ was difficult to analyse in some of the individual spine responses because of noise, the averaged transient yielded a t_1/2_ of ~550 ms, substantially slower than the half-duration of AP-mediated transients recorded in a subset of these spines (n = 8, t_1/2_ of averaged ΔF/F ~100 ms, Fig. 1B bottom). The influence of buffering on the rise time should be less pronounced since the latter mostly reflects the duration of Ca^2+^ entry into the cytoplasm. Indeed, the set of rise times is statistically not different from a previous TPU data set using also DNI-Glutamate and OGB-1 (rise time 76 ± 57 ms, median 60 ms, n = 42, P = 0.19, Mann-Whitney test; data from Bywalez et al., 2015).

**Figure 1.**
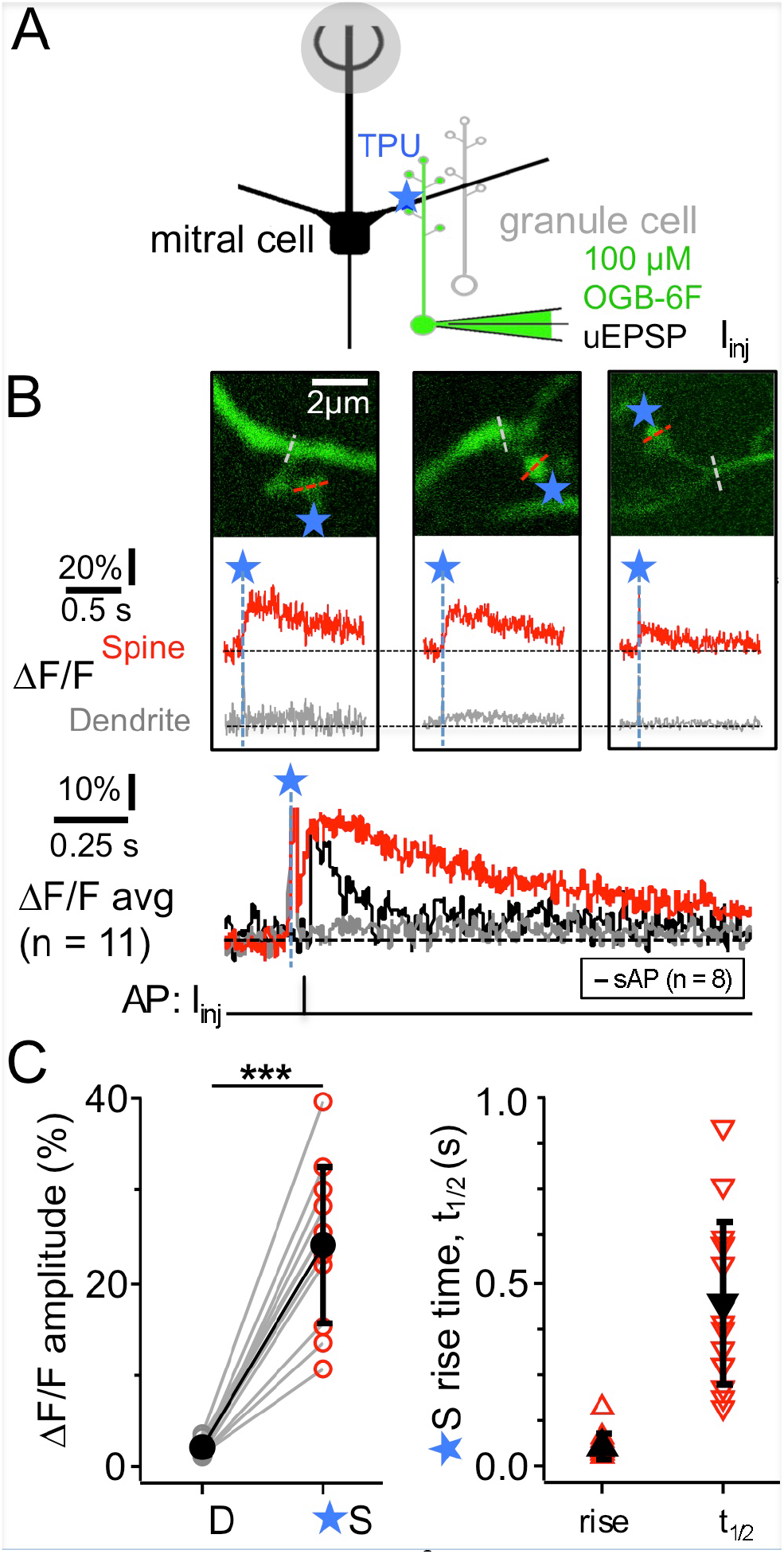
TPU-evoked Ca^2+^ entry into juvenile rat GC spines with low exogenous buffering. Imaging of TPU-evoked Ca^2+^ transients within GC spines with low exogenous buffering (100 μM OGB-6F).. (**A**) Scheme of experiment: Whole cell recording from GC (filled with dye via pipette) and TPU at spine head. The pipette was also used for brief current injections to evoke somatic APs (sAP). (**B**) Top: Two-photon scans (z-projections) of three representative examples of individual spines filled with OBG-6F. Blue stars denote uncaging locations. Red and grey dotted line indicate line scan positions. Middle: Respective averaged fluorescence transients (ΔF/F)_TPU_ that were collected from line scans across the spine heads above (S, red) and the adjacent dendrite at the base of the spine neck (D, grey). Blue dashed lines and star: time point of uncaging. Bottom: (ΔF/F)_TPU_ transients averaged across experiments (Ca^2+^ imaging: n = 11 spines) with the same time axis, spine response in red and dendrite response in grey. The black trace in the Ca^2+^ imaging graph represents the averaged response (ΔF/F)_AP_ to a backpropragating somatically evoked AP (recorded in n = 8 of the 11 spines). (**C**) Cumulative plots of (ΔF/F)_TPU_ amplitudes in dendrite and spine pairs (highly significantly different: P < 0.001, Wilcoxon test), and of rise times and half durations t_1/2_ of (ΔF/F)_TPU_ within the spine heads (mostly not detectable in the dendrites).

### Time course of synaptic spine Na^+^ signals with minimal added exogenous buffering

Postsynaptic Na^+^ signals could report the activity of the Ca^2+^-impermeable GC AMPARs and of spine Na_v_s and TRPC1/4 in a more direct way than Ca^2+^ signals and thus yield additional information on the state of the locally activated GC spine. We performed two-photon Na^+^ imaging using SBFI at a concentration of 1 mM. This is far below both the Na^+^ concentration of 15 mM in the internal solution and the apparent K_D_ of SBFI, so the degree of introduced buffering is negligible (Mondragao et al., 2016; Fig. 2).

**Figure 2.**
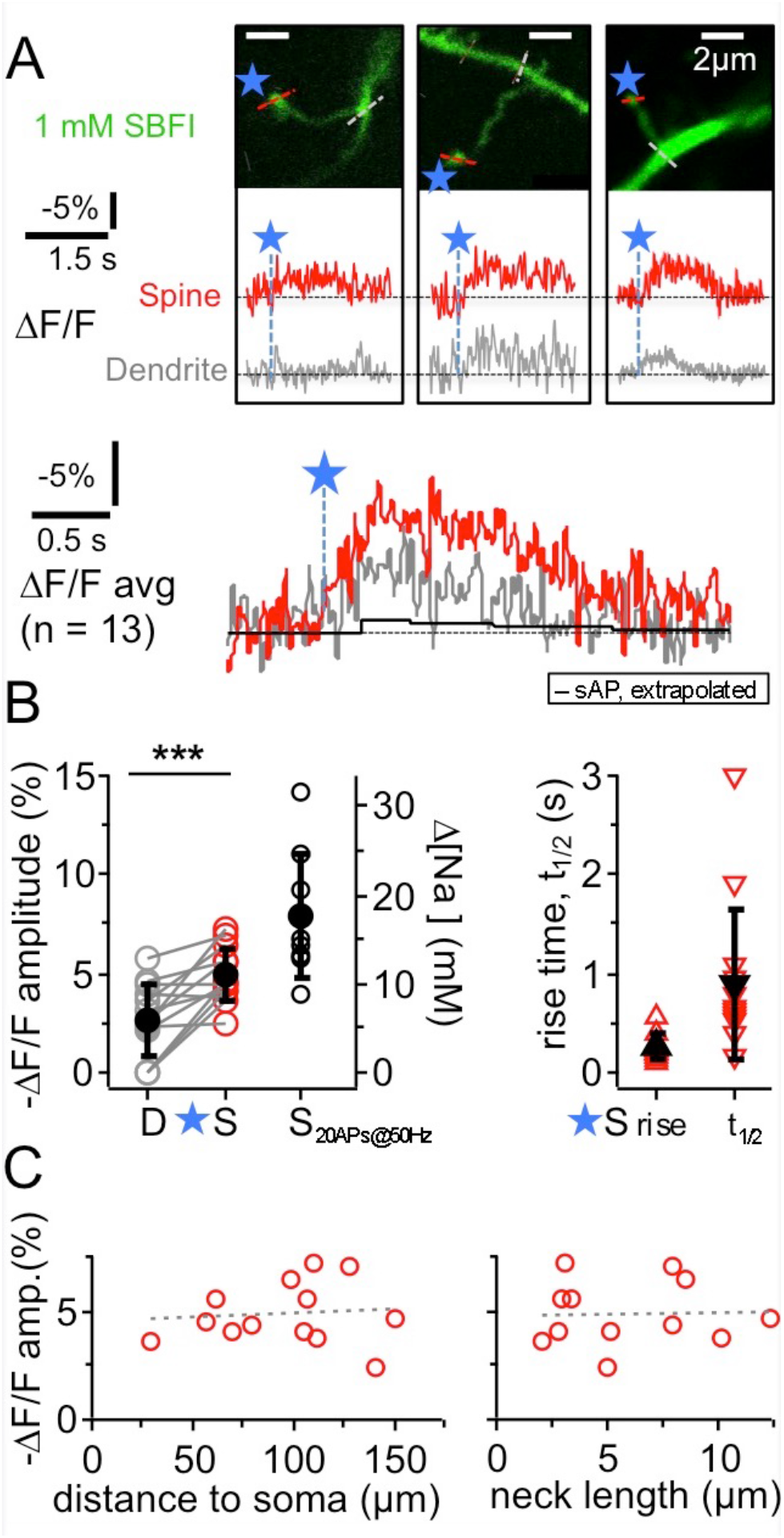
TPU-evoked Na^+^ entry into juvenile rat GC spines. Imaging of TPU-evoked Na^+^ transients within GC spines with low exogenous buffering (1 mM SBFI). Experimental setup as in Fig. 1A. (**A**) See also Fig. 1B. Top: Two-photon z-stacks of three representative examples of individual spines filled with SBFI. Blue stars denote uncaging locations. Red and grey dotted line indicate line scan positions for the leftmost spine. Middle: Respective averaged fluorescence transients (ΔF/F)_TPU_ within the above spines (S, red) and within the adjacent dendrite (at the base of the spine neck, D, grey). Bottom: (ΔF/F)_TPU_ transients averaged across experiments (n = 13 spines) with the same time axis, spine response in red and dendrite response in grey. Note the similar time course of the dendritic response. Black trace: extrapolated response (ΔF/F)_AP_ to a single backpropragating somatically evoked AP (from mean response to 20 APs at 50 Hz, see B. Amplitude (ΔF/F)_AP_ ≈ −0.4 %, t_1/2_ ≈ 1.7 s). (**B**) Cumulative plots. Left panel: (ΔF/F)_TPU_ amplitudes in dendrite and spine pairs (highly significantly different, P < 0.001, Wilcoxon test) and (ΔF/F)_50Hz_ amplitudes in response to a 50 Hz train of 20 APs in a subset of the same spine heads (n = 9). Right axis: Changes in [Na^+^]_i_ concentration (calibration from *Ona-Jodar et al. 2017*). Right panel: Half durations t_1/2_ of (ΔF/F)_TPU_ within the spine heads (t_1/2_ could not be measured in most dendritic transients because of noise). (**C**) (ΔF/F)_TPU_ amplitudes vs. distance of the spine to the soma and spine neck length (means 99 ± 35 μm and 5.9 ± 3.3 μm respectively, n = 12). Dashed lines: Linear fit. No significant correlations.

The ensuing Na^+^ signals following TPU of glutamate at individual spine heads with similar parameters as in the OGB-6F experiments had a mean amplitude of −(ΔF/F)_TPU_ = 4.9 ± 1.4 % in the spine head of juvenile rat GCs (PND 11-18). They were localized to the spine head to some extent but mostly also detectable in the adjacent dendritic shaft (mean amplitude ratio dendrite/spine 0.56 ± 0.38 of spine signal; P < 0.001 vs. spine signal amplitude; n = 13 spines in 11 GCs). Conversion of the spine signal amplitude to absolute changes in [Na^+^]_i_ (Rose et al., 1999; Ona-Jodar et al., 2017) yielded a mean increase Δ[Na^+^]_i_ by ~10 mM. The average rise time was 250 ± 130 ms and the half duration t_1/2_ = 890 ± 770 ms in the spines, including frequently observed plateau phases. Individual (ΔF/F)_TPU_ signals in dendritic shafts were usually too noisy for kinetic analysis.

As for TPU-evoked Ca^2+^ transients (Bywalez et al., 2015), there were no significant correlations between spine (ΔF/F)_TPU_ amplitudes and distance to soma or spine neck length (Fig. 2C), and also not between the amplitude ratio of spine/dendrite and spine neck length (not shown).

Again, we averaged data across all spine/dendrite pairs (Fig. 2A bottom). The averaged spine signal showed an initial plateau phase of ~600 ms, and the averaged dendrite signal mirrored the kinetics of the spine signal, which is expected because of the fast diffusion of Na^+^ into the dendrite (Mondragao et al., 2016). Still, these TPU-evoked Na^+^ signals are very slow in view of the overall fast diffusion of Na^+^ and also compared to recent data from synaptic Na^+^ signals in hippocampal pyramidal neuron spines (their t_1/2_ ~20 ms; Miyazaki and Ross, 2017), and therefore are best explained by a persistent influx of Na^+^ (see discussion). The TPU-evoked spine Na^+^ signals are also very large as compared to the Na^+^ influx induced by single backpropagating APs. The latter was on the order of 0.4 mM (as extrapolated from train stimulation of a subset of the same spines with 20 APs at 50 Hz: (ΔF/F)_50Hz_ = −7.9 ± 3.1 % n = 9, Fig. 2B; see Fig. 5 in Ona-Jodar et al., 2017 for train responses).

From these experiments we conclude that the time course of asynchronous components of GABA release triggered by unitary activation (Lage-Rupprecht et al., 2020) matches well with substantial and prolonged elevation of postsynaptic Na^+^ and Ca^2+^ concentrations in the GC spine. Late release therefore might result from local processing following unitary inputs to the reciprocal spine (see Discussion).

### Postsynaptic GC spine Ca^2+^ signals in adult mice

As shown above, both postsynaptic Ca^2+^ and Na^+^ signalling in juvenile rat GC spines is likely to persist for several 100 ms. Moreover, we noticed that in spite of similar endogenous Ca^2+^ dynamics (with regard to both buffering capacity and extrusion: Egger and Stroh, 2009), postsynaptic spine Ca^2+^ transients in adult mice evolved yet more sluggishly. Here we analysed the kinetics and response probability of postsynaptic GC spine Ca^2+^ signals in response to glomerular stimulation from an earlier data set that was recorded with two-photon fluorescence imaging in acute bulb slices from WT animals or from animals with partial GC GluN1 or GluA2 deletions via viral transfection (PND 36-66; dye 100 μM OGB-1; Abraham et al. 2010).

Postsynaptic Ca^2+^ transients in WT adult mouse GC spines were also strictly localized to spine heads and occurred with a rather low probability upon glomerular stimulation, even though compound EPSPs were readily recorded at the soma (see Fig. 3B), rendering the set of recorded individual responses rather small (estimated response probability P_r_ 0.10 ± 0.06, n = 13 spines; Fig. 3A, Fig. 4A-C, see Methods). While some responses rose rather quickly (e.g. top left response in Fig. 3B, distribution in Fig. 4B, C), in most cases the peak amplitude of these (ΔF/F)_syn_ signals occurred several 100 ms later (average time to peak (TTP) 420 ± 440 ms, beginning with 20 ms, n = 14 events; 2 events with peak beyond scan time of 1000 ms; if these are included with the end of scan as peak time: TTP_min_ = 540 ± 510 ms). TTP was uncorrelated to peak amplitude (r = 0.32, P = 0.11, n = 14). Such slowly rising signals were never observed in the adjacent dendrite. We also recorded responses to single backpropagating APs (ΔF/F)_sAP_ in 9 of the 13 spines, with mean rise times of 15 ± 4 ms (see also Stroh et al 2012).

**Figure 3.**
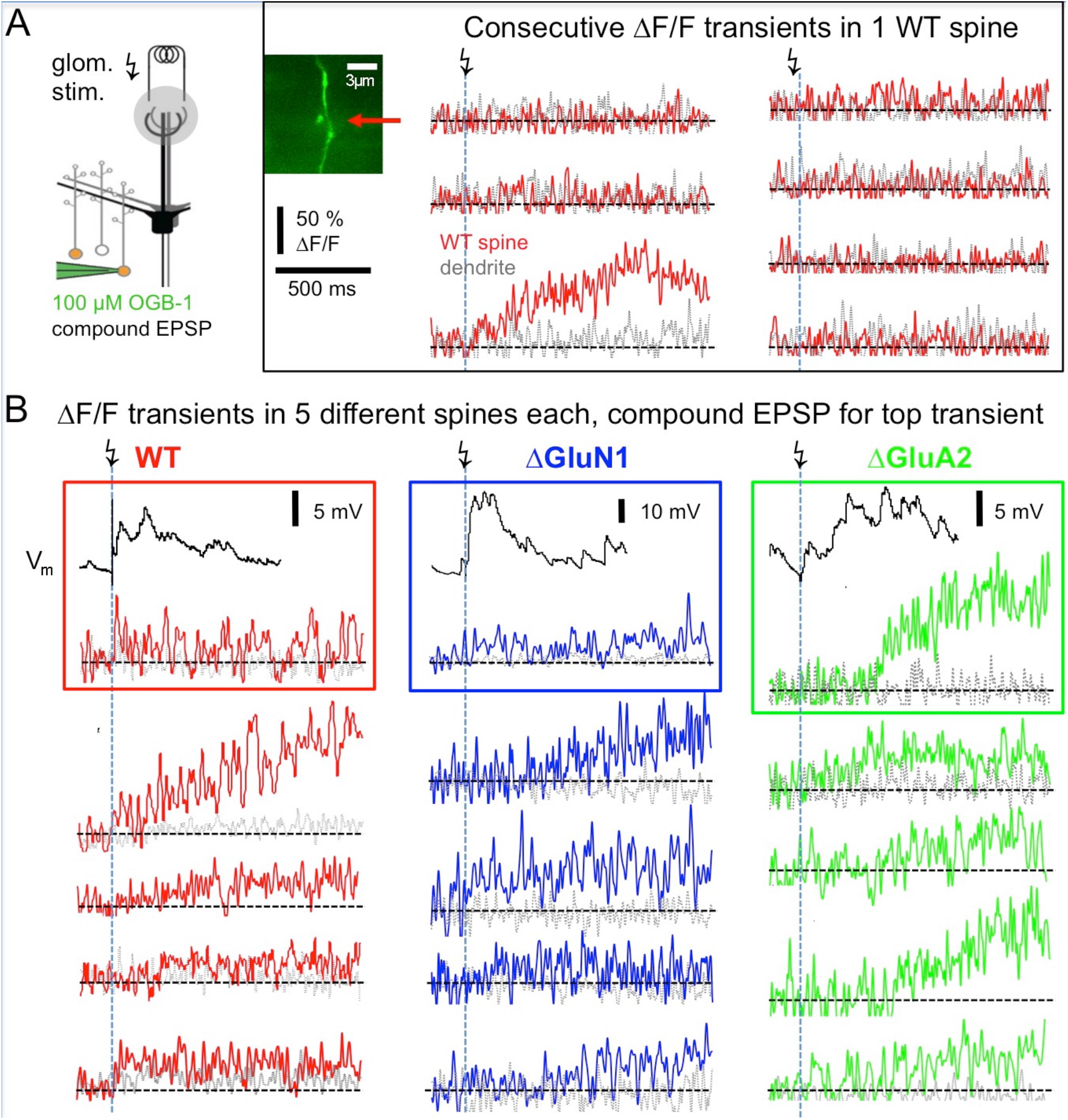
Postsynaptic responses in adult WT, ΔGluA2 and ΔGluN1 GC spines. (**A**) Left: Scheme of experiment (modified from *Abraham et al. 2010*, their Fig. 2). Electrical glomerular stimulation and whole-cell recording of the resulting compound EPSP from a GC, including two-photon imaging of a responding spine and its adjacent dendritic shaft. Genetically modified GCs (ΔGluA2 and ΔGluN1) were identified by the expression of the fluorescent construct Kusabira-Orange. Middle: Scan of a WT spine and dendrite. Right: Line scans through the spine and dendrite during consecutive glomerular stimulations. Only one response occurred (3^rd^ trace), that shows a slowly evolving Ca^2+^ transient confined to the spine head. (**B**) Single responses from other spines, with each trace imaged in a different spine. Same scales for ΔF/F and time as in (A). Top transients with their associated compound EPSP recordings. Left (red): WT GCs. Middle (blue): ΔGluN1 GCs. Right (green): ΔGluA2 GCs. Note the larger size and yet slower evolution compared to WT and ΔGluN1.

**Figure 4.**
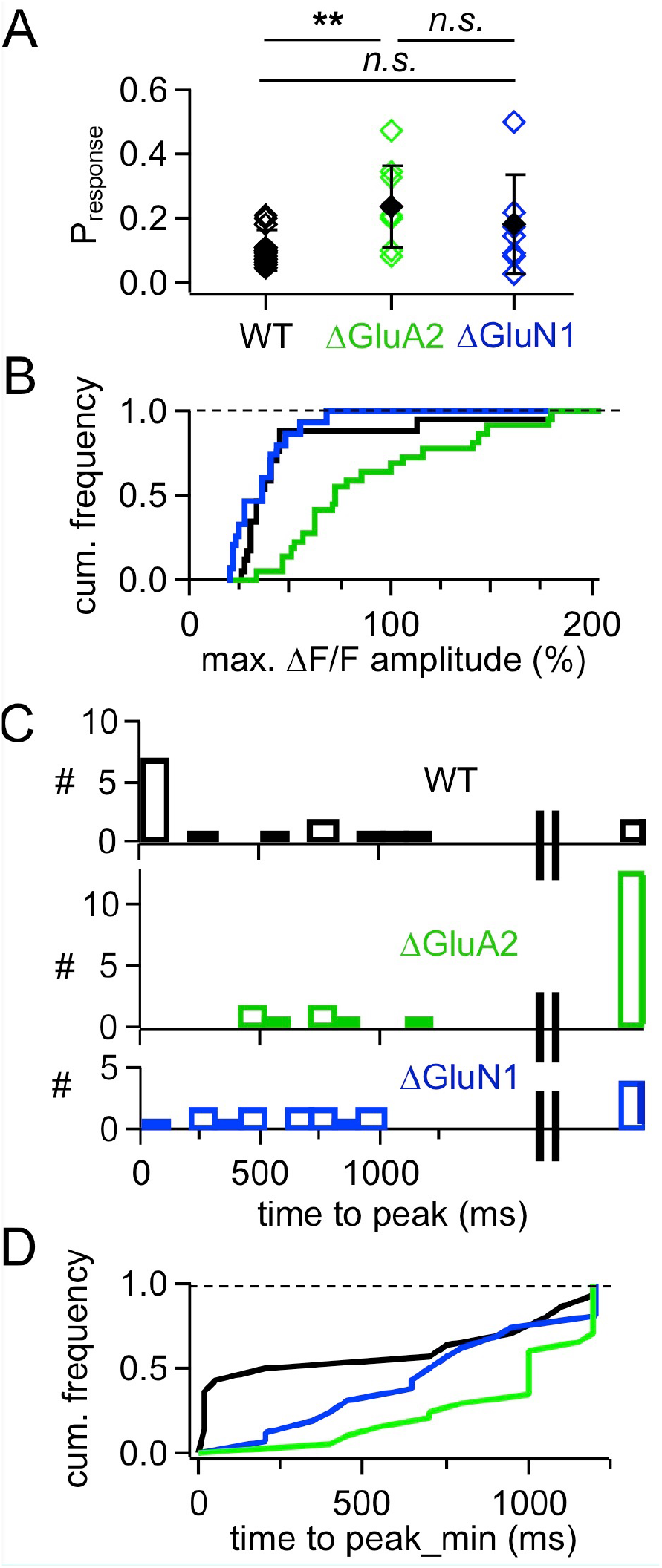
Cumulative analyses of responses in adult mouse WT, ΔGluA2 and ΔGluN1 GCs. (**A**) Estimated likelihood of observing a (ΔF/F)_syn_ response upon glomerular stimulation (‘response probability’, calculated as number of responses divided by number of stimulations). Pairwise comparison (Mann-Whitney test) and Bonferroni: P_r_ WT vs ΔGluA2: P = 0.0018, ΔGluA2 vs ΔGluN1: P = 0.42, WT vs ΔGluN1: P = 0.51 (**B**) Frequency distribution of (ΔF/F)_syn_ response amplitudes for the three GC groups. WT is different from ΔGluA2, but not from ΔGluN1 (P < 0.001 and P = 0.16, Kolmogorov-Smirnov). (**C**) Distributions of (ΔF/F)_syn_ response time to peak for the three GC groups. The rightmost bar shows the responses where the peak might not yet have been reached within the scanning time window (up to 1200 ms post stimulation). (**D**) Frequency distribution of (ΔF/F)_syn_ response time to peak for the three GC groups, with peaks possibly beyond scan window set to 1200 ms (minimal time to peak). WT is different from ΔGluA2, but not from ΔGluN1 (P = 0.02 and P = 0.09, Kolmogorov-Smirnov test).

This slow rise was unrelated to Ca^2+^ entry via NMDARs, since similarly slow signals occurred in ΔGluN1 GC spines with reduced Ca^2+^ entry (Fig. 3B and 4A-C; ΔGluN1 TTP 580 ± 260 ms, beginning with 200 ms, n = 13 events; 4 events with peak beyond scan, TTP_min_ = 740 ± 350 ms; no significant difference to WT TTP_min_: Kolmogorov-Smirnov test P = 0.09). The probability to observe such events in ΔGluN1 GCs (P_r_ = 0.18 ± 0.15, n = 7 spines) was also not significantly different from WT.

Additional Ca^2+^ entry due to GluA2 deletion, however, resulted in larger ΔF/Fs that rose even more slowly than in WT spines (Fig. 3B and 4A-C; ΔGluA2 TTP 680 ± 250 ms, beginning with 400 ms, n = 7 events; 13 events with peak beyond scan; all events: TTP_min_ = 960 ± 270 ms: Kolmogorov-Smirnov test P < 0.02 vs WT). The probability of such events was significantly higher than in WT (P_r_ = 0.24 ± 0.13, n = 8 spines, P = 0.006), possibly due to their improved detectability because of larger ΔF/F amplitudes. Such a slow evolution in postsynaptic Ca^2+^ should also be reflected in the strength and time course of reciprocal GABA release. Indeed, it was shown previously in recordings of mitral cell hyperpolarizing potentials following a train of 20 APs at 50 Hz (a stimulation that efficiently activates reciprocal release from GCs; Fig. 4 in Abraham et al. 2010), that ΔGluA2 resulted in a significantly stronger and almost two times longer mitral cell recurrent inhibition compared to WT. The relevant time scales match our time to peak data including the late peaks of (ΔF/F)_syn_ reported above (ΔGluA2: half duration t_1/2_ ~ 1150 ms; WT: t_1/2_ ~ 600 ms). Conversely, ΔGluN1 did not exert a significant influence on the duration of recurrent inhibition and only a mild reduction on its amplitude, in line with the lack of a significant effect on TTP of (ΔF/F)_syn_ described above (Supplementary Information in Abraham et al. 2010).

## Discussion

### Relation between Ca^2+^ and Na^+^ transients in the GC spine head and asynchronous release

Asynchronous release – i.e. release that happens later than the fast coupling of HVA presynaptic Ca^2+^ currents to the release machinery (e.g. Kaeser and Regehr, 2014) - is a phenomenon known from many central synapses. It is often observed at repetitively stimulated synapses (Wen et al., 2013), which would also hold true for the classical DDI protocol, where voltage-clamped mitral cells are being depolarized for 20-50 ms and thus ongoing release of glutamate from mitral cells is likely to happen over dozens of ms and subsequent asynchronous release of GABA has been documented by many groups (see Introduction). Thus it is at first surprising that local, unitary-like stimulation of GC spines by TPU would suffice to elicit asynchronous release, which we have recently documented (Lage-Rupprecht et al., 2020). However, the temporal extent of this asynchronous release was shorter than in the classical DDI experiments (maximal extent of ~500 ms vs > 1 s) and therefore there might be additional mechanisms involved whenever GCs are activated more strongly.

The NMDAR-mediated Ca^2+^ current into juvenile rat GC spines is expected to recede within less than 200 ms (e.g. Egger et al., 2005; Hestrin et al., 1990). It is thus on its own unlikely to mediate substantial asynchronous release far beyond the first 100 ms, even though the slow extrusion of Ca^2+^ from the cytoplasm may contribute to delayed release (Egger and Stroh, 2009). To further unravel signalling downstream of the NMDAR and Na_v_ activation during the local spine spike, we investigated the time course of postsynaptic Na^+^ and Ca^2+^ elevations with minimal exogenous buffering. Both ion species showed prolonged elevations for durations well compatible with asynchronous output.

Previously, somatic AP-mediated and postsynaptic Ca^2+^ transients recorded with 100 μM OGB-1 were observed to decay with roughly equal half durations of 600 ms (except for the subpopulation of ‘slow spines’ featuring transients with t_1/2_ > 1.5 s, which made up one third; Egger et al. 2005; Bywalez et al. 2015). Whereas here TPU-evoked transients were always substantially longer than AP-evoked transients in the same spine, on the order of 500 ms vs 100 ms, dissociating physiological postsynaptic Ca^2+^ dynamics from exogenous buffer effects. Interestingly, ‘slow spines’ were not observed here, also not in an additional set of n = 12 spine responses that was excluded from the analysis because of inadvertently longer uncaging intervals. This observation might be explained by the existence of a Ca^2+^-dependent extrusion mechanism, that is activated only by high levels of [Ca^2+^]_i_ which are buffered away in the presence of 100 μM OGB-1 (such as Ca^2+^-ATPases, Na^+^/Ca^2+^ exchangers or mitochondrial Ca^2+^ uniporters, e.g. Sabatini et al., 2002; Chamberland et al., 2019).

In particular, there was a substantial and long-lasting postsynaptic elevation of Na^+^. This detected Δ[Na^+^]_i_ is ~ 20fold higher than what could be extrapolated for a single backpropagating GC AP from responses to train stimulation (~10 mM vs. ~0.4 mM, see Fig. 2A). Thus there must be substantial postsynaptic Na^+^ entry on top of local Na_v_ activation, in line with earlier reports of Na^+^ signals in response to suprathreshold synaptic stimulation in hippocampal pyramidal neurons (Rose and Konnerth, 2001). In GC spines, the average time course of Δ[Na^+^]_i_ showed a plateau-like phase of >500 ms, very much unlike recent observations of single synaptic Na^+^ transients in spines of hippocampal pyramidal neurons which decayed within 20 ms – but were of a similar magnitude (~5 mM; Miyazaki and Ross, 2017). While the slow decay of the GC spine Δ[Na^+^]_i_ might be explained to some extent by the diffusive barrier provided by the high neck resistance (predicted as ≥1 GΩ, Bywalez et al., 2015), the origin of the [Na^+^]_i_ plateau requires a persistent Na^+^ entry that outlasts AMPAR/NMDAR activation. It thus might indeed be related to extended local TRPC1/4 activation downstream of NMDAR activation, since there was no global plateau current in ΔGluN1 GCs (Stroh et al., 2012).

Alternatively or in addition, the [Na^+^]_i_ plateau might correspond to a local UP state in the GC spine which could cause TRPC1/4 activation and thus ongoing local influx of Ca^2+^ sufficient to trigger recurrent release. This influx should also happen close to the release machinery, since buffering of GC Ca^2+^ by EGTA had no effect on asynchronous release (Isaacson, 2001). Local UP states might not be evident in GC somatic membrane potential recordings due to substantial filtering by the large spine neck resistance. Increased [Na^+^]_i_ within the observed regime might also provide positive feedback to NMDARs via an upregulation of NMDAR Ca^2+^ currents by the Src kinase (Yu and Salter, 1997, 1998). Further experiments are required to unravel such interactions.

The large size of Δ[Na^+^]_i_ amplitudes in the adjacent dendrite is unexpected, since Na^+^ diffusion from the spine into the dendrite should result in a substantial drop in concentration (e.g. Miyazaki & Ross 2017) and GC spine necks are particularly long (~ 6 μm in our sample, Fig 2C). Possible explanations for this observation include the large volume of GC spine heads (similar radius as the dendrite, Egger & Stroh 2009), weak Na^+^ extrusion from the spine head and neck, or that the source of the synaptically triggered persistent Na^+^ entry mechanism postulated above is present and activated also within the dendritic shaft. In any case, dendritic voltage-gated Na_v_ channels (e.g. Egger et al., 2003; Nunes and Kuner, 2018) are unlikely to contribute to this signal since we have shown previously that the postsynaptic spine depolarization undergoes strong attenuation and thus both dendritic Na_v_ and Ca_v_ channels will not become activated (Bywalez et al., 2015) - unless several spines receive coincident inputs (Mülller and Egger, 2020).

### Postsynaptic Ca^2+^ signals in adult mouse versus juvenile rat

Endogenous Ca^2+^ buffering and extrusion, postsynaptic ΔCa^2+^ amplitudes and synaptic AP-evoked Ca^2+^ signals and long-lasting depolarizations are similar in adult mouse GCs as in juvenile rat GCs (Egger et al., 2005; Egger, 2008; Egger and Stroh, 2009; Abraham et al., 2010; Stroh et al., 2012). However, a more detailed analysis of the synaptic responses showed also two striking differences. First, we observed a strongly reduced release probability P_r_ (0.1 vs. 0.5). This effect might be due to maturation of the bulbar network, since in rats the strength of dendrodendritic inhibition was reported to decline steeply between PND15 and PND20 (Dietz et al., 2011). While this P_r_ value is no more than an estimate due to the small number of recorded responses and substantial noise in some experiments, we observed similar values also for recordings from ΔGluN1 and ΔGluA2 GCs, with the slight increase for ΔGluA2 possibly explained by the improved detectability of signals in these cells. Second, yet more strikingly, we observed a broad variability in signal onset and rise. In all three types of GCs, there was a subset of signals with an apparently delayed onset and a very slow evolution within hundreds of milliseconds up to seconds. These features were unchanged between WT and ΔGluN1, so NMDARs are not required to generate such responses. Rather, the ΔGluA2 GCs showed an increased fraction of such slow signals, indicating that Ca^2+^-permeable AMPARs can also feed into this mechanism. This effect could be mediated perhaps by enhancing postsynaptic Ca^2+^ induced Ca^2+^ release (CICR) that we have previously shown to also occur in rat GC spines (Egger et al., 2005). Such a Ca^2+^-dependent mechanism might also be supported by slow extrusion (Egger and Stroh, 2009). Together with the current observations these data are consistent with voltage-gated Ca^2+^ channels or Ca^2+^-permeable AMPARs triggering CICR, rather than NMDARs, as was also observed in other neuron types (e.g. Chavez et al., 2006; Plotkin et al., 2013). In any case, the respective mechanism is also likely to undergo developmental upregulation since a delayed and extended postsynaptic Ca^2+^ rise was not observed in young rat GC spines.

Intriguingly, apparently slowly rising signals in GC spines may also be of presynaptic origin, e.g. due to late firing of principal neurons in response to glomerular stimulation (Kapoor and Urban, 2006; Giridhar and Urban, 2012; Gire et al., 2012). In our experiments, this source might also contribute in the wake of glomerular stimulation; a correlation with late EPSPs is difficult to test because of the high spontaneous activity and low number of responses in our recordings. However, such delayed presynaptic activity is probably not a main source of slowly rising signals in our data set, since a enhancement of presynaptic signal contributions specifically in ΔGluA2 GCs appears rather unlikely.

As already implied by this possibility of delayed presynaptic inputs, slow signals in olfactory bulb networks are not restricted to GC-mediated recurrent inhibition; representations of olfactory stimuli in general are known to evolve over long time scales of hundreds to thousands of ms (Friedrich and Laurent, 2001; Uchida and Mainen, 2003; Abraham et al., 2004, 2010; Rinberg et al., 2006, Gschwend et al., 2015). Aside from the original notion that these time scales are required for decorrelation of principal neuron activity, persistent representations might also be involved in the formation of odor after-images (Patterson et al., 2013).

With regard to further functions of slow signals, they are at first glance unlikely to play a direct role during odor discrimination or background segregation, since these discriminations usually occur within considerably less than 500 ms, even for difficult mixtures and/or many components (Abraham et al., 2004, 2010; Bhattacharjee et al., 2019; Kepecs et al., 2005; Rokni et al., 2014). Rather, slow signals may be involved in learning and plasticity, also during learning of the mixture discrimination task (Abraham et al., 2004, 2010; Gschwend et al., 2015).

Aberrant slow signals due to pathological changes (extended or reduced asynchronous release) would thus be expected to disrupt plasticity induction. Indeed, several pathologies such as Alzheimer’s disease have been associated with enhanced asynchronous release (reviewed in Kaeser and Regehr 2014), for example at the neuromuscular junction, and interestingly also for fast-spiking interneurons in epileptic foci in both human and rat (Jiang et al., 2012). However, so far no loss of function has been observed for the ΔGluA2 modification. Pathological or other modulations of asynchronous release from granule cells might also influence both slow and fast network oscillations, i.e. the respiration coupled θ and probably more importantly γ rhythm that GCs have been associated with (e.g. Fukunaga et al., 2014). Such interactions between asynchronous release and rhythmic activity have been demonstrated in other GABAergic interneurons, including hippocampal cholecystokinin-positive basket interneurons that are crucially involved in generation of the hippocampal theta rhythm (e.g. Hefft and Jonas, 2005), and in cortical parvalbumin-positive GABAergic neurons that power γ oscillations (e.g. Traub et al., 1998). Interestingly, increased asynchronous release of GABA might reduce the ability of these PV+ neurons to sustain γ and has been proposed as one possible mechanism for uncoupling in schizophrenia (Volman et al., 2011).

In conclusion, we find that several mechanisms such as delayed and slowly evolving excitation, slow removal of Ca^2+^ and perhaps extended local postsynaptic depolarization as indicated by the persistent elevation of Na^+^ may feed into asynchronous GC spine output. Since on the other hand there is also a fast, synchronous component of reciprocal release (Halabisky et al., 2000; Lage-Rupprecht et al., 2020), GC spines are obviously capable of parallel processing on multiple time scales, a property that appears to be further refined with maturation.

## Author contributions

Rat granule cell Ca^2+^ imaging was performed by VLR, rat granule cell Na^+^ imaging by TOJ, mouse viral injections by NA and mouse granule cell Ca^2+^ imaging by VE. VLR, TOJ and VE analyzed data. VE, NA and CR designed research. VE wrote the manuscript. All co-authors except for TOJ contributed to editing the manuscript.

## Acknowledgements

We wish to thank A. Pietryga-Krieger for expert technical assistance, Dr. N. Gerkau (Heinrich Heine University Düsseldorf) for initial help with Na^+^ imaging, and B. Rozsa for DNI-glutamate. The instrumentation used in this study was funded by the BMBF (FKZ 01GQ1502; VE) with additional equipment funding by DFG-SFB 870 (EG135/5-1) and GSN-LMU.

## References

Abraham NM, Egger V, Shimshek DR, Renden R, Fukunaga I, Sprengel R, Seeburg PH, Klugmann M, Margrie TW, Schaefer AT, Kuner T (2010) Synaptic Inhibition in the Olfactory Bulb Accelerates Odor Discrimination in Mice. Neuron 65:399–411.

Abraham NM, Spors H, Carleton A, Margrie TW, Kuner T, Schaefer AT (2004) Maintaining accuracy at the expense of speed: stimulus similarity defines odor discrimination time in mice. Neuron 44:865–876.

Bhattacharjee AS, Konakamchi S, Turaev D, Vincis R, Nunes D, Dingankar AA, Spors H, Carleton A, Kuner T, Abraham NM (2019) Similarity and Strength of Glomerular Odor Representations Define a Neural Metric of Sniff-Invariant Discrimination Time. Cell Reports 28:2966–2978.e2965.

Buonviso N, Amat C, Litaudon P, Roux S, Royet J-P, Farget V, Sicard G (2003) Rhythm sequence through the olfactory bulb layers during the time window of a respiratory cycle. European Journal of Neuroscience 17:1811–1819.

Bywalez Wolfgang G, Patirniche D, Rupprecht V, Stemmler M, Herz Andreas VM, Pálfi D, Rózsa B, Egger V (2015) Local Postsynaptic Voltage-Gated Sodium Channel Activation in Dendritic Spines of Olfactory Bulb Granule Cells. Neuron 85:590–601.

Chamberland S, Moratalla AZ, Topolnik L (2019) Calcium extrusion mechanisms in dendrites of mouse hippocampal CA1 inhibitory interneurons. Cell Calcium 77:49–57. doi: 10.1016/j.ceca.2018.12.002. Epub 2018 Dec 4.

Chávez AE, Singer JH, Diamond JS (2006) Fast neurotransmitter release triggered by Ca influx through AMPA-type glutamate receptors. Nature 443:705–708.

Chen WR, Xiong W, Shepherd GM (2000) Analysis of relations between NMDA receptors and GABA release at olfactory bulb reciprocal synapses. Neuron 25:625–633.

Dietz SB, Markopoulos F, Murthy VN (2011) Postnatal development of dendrodendritic inhibition in the Mammalian olfactory bulb. Front Cell Neurosci 5:10.

Egger V (2008) Synaptic sodium spikes trigger long-lasting depolarizations and slow calcium entry in rat olfactory bulb granule cells. European Journal of Neuroscience 27:2066–2075.

Egger V, Stroh O (2009) Calcium buffering in rodent olfactory bulb granule cells and mitral cells. The Journal of physiology 587:4467–4479.

Egger V, Svoboda K, Mainen ZF (2003) Mechanisms of Lateral Inhibition in the Olfactory Bulb: Efficiency and Modulation of Spike-Evoked Calcium Influx into Granule Cells. The Journal of Neuroscience 23:7551–7558.

Egger V, Svoboda K, Mainen ZF (2005) Dendrodendritic Synaptic Signals in Olfactory Bulb Granule Cells: Local Spine Boost and Global Low-Threshold Spike. The Journal of Neuroscience 25:3521–3530.

Friedrich RW, Laurent G (2001) Dynamic optimization of odor representations by slow temporal patterning of mitral cell activity. Science 291:889–894.

Fukunaga I, Herb JT, Kollo M, Boyden ES, Schaefer AT (2014) Independent control of gamma and theta activity by distinct interneuron networks in the olfactory bulb. Nat Neurosci 17:1208–1216.

Gire DH, Franks KM, Zak JD, Tanaka KF, Whitesell JD, Mulligan AA, Hen R, Schoppa NE (2012) Mitral cells in the olfactory bulb are mainly excited through a multistep signaling path. J Neurosci 32:2964–2975.

Giridhar S, Urban NN (2012) Mechanisms and benefits of granule cell latency coding in the mouse olfactory bulb. Frontiers in neural circuits 6:40–40.

Gschwend O, Abraham NM, Lagier S, Begnaud F, Rodriguez I, Carleton A (2015) Neuronal pattern separation in the olfactory bulb improves odor discrimination learning. Nature neuroscience 18:1474–1482.

Halabisky B, Friedman D, Radojicic M, Strowbridge BW (2000) Calcium influx through NMDA receptors directly evokes GABA release in olfactory bulb granule cells. The Journal of neuroscience : the official journal of the Society for Neuroscience 20:5124–5134.

Hall BJ, Delaney KR (2002) Contribution of a calcium-activated non-specific conductance to NMDA receptor-mediated synaptic potentials in granule cells of the frog olfactory bulb. The Journal of physiology 543:819–834.

Hefft S, Jonas P (2005) Asynchronous GABA release generates long-lasting inhibition at a hippocampal interneuron–principal neuron synapse. Nature Neuroscience 8:1319–1328.

Hestrin S, Sah P, Nicoll RA (1990) Mechanisms generating the time course of dual component excitatory synaptic currents recorded in hippocampal slices. Neuron 5:247–253.

Isaacson JS (2001) Mechanisms governing dendritic gamma-aminobutyric acid (GABA) release in the rat olfactory bulb. Proc Natl Acad Sci U S A 98:337–342.

Isaacson JS, Strowbridge BW (1998) Olfactory Reciprocal Synapses: Dendritic Signaling in the CNS. Neuron 20:749–761.

Jardemark K, Nilsson M, Muyderman H, Jacobson I (1997) Ca^2+^ ion permeability properties of (R,S) alpha-amino-3-hydroxy-5-methyl-4-isoxazolepropionate (AMPA) receptors in isolated interneurons from the olfactory bulb of the rat. J Neurophysiol 77:702–708.

Jiang M, Zhu J, Liu Y, Yang M, Tian C, Jiang S, Wang Y, Guo H, Wang K, Shu Y (2012) Enhancement of asynchronous release from fast-spiking interneuron in human and rat epileptic neocortex. PLoS biology 10:e1001324–e1001324.

Kaeser PS, Regehr WG (2014) Molecular mechanisms for synchronous, asynchronous, and spontaneous neurotransmitter release. Annual review of physiology 76:333–363.

Kapoor V, Urban NN (2006) Glomerulus-specific, long-latency activity in the olfactory bulb granule cell network. The Journal of neuroscience : the official journal of the Society for Neuroscience 26:11709–11719.

Kepecs A, Uchida N, Mainen ZF (2005) The Sniff as a Unit of Olfactory Processing. Chemical Senses 31:167–179.

Lage-Rupprecht V, Zhou L, Bianchini G, Aghvami SS, Rózsa B, Sassoé-Pognetto M, Egger V (2020) Presynaptic NMDA receptors cooperate with local action potentials to implement activity-dependent GABA release from the reciprocal olfactory bulb granule cell spine. bioRxiv:440198.

Lagier S, Carleton A, Lledo P-M (2004) Interplay between local GABAergic interneurons and relay neurons generates gamma oscillations in the rat olfactory bulb. The Journal of neuroscience : the official journal of the Society for Neuroscience 24:4382–4392.

Miyazaki K, Ross WN (2017) Sodium Dynamics in Pyramidal Neuron Dendritic Spines: Synaptically Evoked Entry Predominantly through AMPA Receptors and Removal by Diffusion. The Journal of neuroscience : the official journal of the Society for Neuroscience 37:9964–9976.

Mondragão MA, Schmidt H, Kleinhans C, Langer J, Kafitz KW, Rose CR (2016) Extrusion versus diffusion: mechanisms for recovery from sodium loads in mouse CA1 pyramidal neurons. The Journal of physiology 594:5507–5527.

Mueller M, Egger V (2020) Dendritic integration in olfactory bulb granule cells: Threshold for lateral inhibition and role of active conductances upon simultaneous activation. PLoS Biol, in press. bioRxiv:901397. doi.org/10.1101/2020.01.10.90139

Nunes D, Kuner T (2018) Axonal sodium channel NaV1.2 drives granule cell dendritic GABA release and rapid odor discrimination. PLoS Biol, 16(8), e2003816. doi:10.1371/journal.pbio.2003816

Nusser Z, Kay LM, Laurent G, Homanics GE, Mody I (2001) Disruption of GABAA Receptors on GABAergic Interneurons Leads to Increased Oscillatory Power in the Olfactory Bulb Network. Journal of Neurophysiology 86:2823–2833.

Ona-Jodar T, Gerkau NJ, Sara Aghvami S, Rose CR, Egger V (2017) Two-Photon Na(+) Imaging Reports Somatically Evoked Action Potentials in Rat Olfactory Bulb Mitral and Granule Cell Neurites. Frontiers in cellular neuroscience 11:50–50.

Osinski BL, Kay LM (2016) Granule cell excitability regulates gamma and beta oscillations in a model of the olfactory bulb dendrodendritic microcircuit. Journal of Neurophysiology 116:522–539.

Patterson MA, Lagier S, Carleton A (2013) Odor representations in the olfactory bulb evolve after the first breath and persist as an odor afterimage. Proc Natl Acad Sci U S A 110:E3340–3349.

Plotkin JL, Shen W, Rafalovich I, Sebel LE, Day M, Chan CS, Surmeier DJ (2013) Regulation of dendritic calcium release in striatal spiny projection neurons. J Neurophysiol 110:2325–2336.

Rall W, Shepherd GM (1968) Theoretical reconstruction of field potentials and dendrodendritic synaptic interactions in olfactory bulb. Journal of Neurophysiology 31:884–915.

Rinberg D, Koulakov A, Gelperin A (2006) Sparse odor coding in awake behaving mice. J Neurosci 26:8857–8865.

Rokni D, Hemmelder V, Kapoor V, Murthy VN (2014) An olfactory cocktail party: figure-ground segregation of odorants in rodents. Nature Neuroscience 17:1225–1232.

Rose CR, Konnerth A (2001) NMDA receptor-mediated Na^2+^ signals in spines and dendrites. J Neurosci 21:4207–4214.

Rose CR, Kovalchuk Y, Eilers J, Konnerth A (1999) Two-photon Na^2+^ imaging in spines and fine dendrites of central neurons. Pflügers Archiv 439:201–207.

Sabatini B, Oertner TG, Svoboda K (2002) The life cycle of Ca^2+^ ions in dendritic spines. Neuron, Vol. 33, 439–452.

Schoppa NE, Kinzie JM, Sahara Y, Segerson TP, Westbrook GL (1998) Dendrodendritic inhibition in the olfactory bulb is driven by NMDA receptors. The Journal of neuroscience : the official journal of the Society for Neuroscience 18:6790–6802.

Schoppa NE, Westbrook GL (1999) Regulation of synaptic timing in the olfactory bulb by an A-type potassium current. Nature Neuroscience 2:1106–1113.

Schoppa NE, Westbrook GL (2001) Glomerulus-Specific Synchronization of Mitral Cells in the Olfactory Bulb. Neuron 31:639–651.

Shepherd GM (2004) Olfactory bulb. The Synaptic Organization of the Brain.

Stroh O, Freichel M, Kretz O, Birnbaumer L, Hartmann J, Egger V (2012) NMDA receptor-dependent synaptic activation of TRPC channels in olfactory bulb granule cells. The Journal of neuroscience 32:5737–5746.

Tang W, Ehrlich I, Wolff SB, Michalski AM, Wolfl S, Hasan MT, Luthi A, Sprengel R (2009) Faithful expression of multiple proteins via 2A-peptide self-processing: a versatile and reliable method for manipulating brain circuits. J Neurosci 29:8621–8629.

Tran V, Park MCH, Stricker C (2018) An improved measurement of the Ca(2+)-binding affinity of fluorescent Ca(2+) indicators. Cell Calcium 71:86–94.

Traub RD, Spruston N, Soltesz I, Konnerth A, Whittington MA, Jefferys GR (1998) Gamma-frequency oscillations: a neuronal population phenomenon, regulated by synaptic and intrinsic cellular processes, and inducing synaptic plasticity. Prog Neurobiol 55:563–575.

Uchida N, Mainen ZF (2003) Speed and accuracy of olfactory discrimination in the rat. Nature Neuroscience 6:1224–1229.

Volman V, Behrens MM, Sejnowski TJ (2011) Downregulation of parvalbumin at cortical GABA synapses reduces network gamma oscillatory activity. The Journal of neuroscience : the official journal of the Society for Neuroscience 31:18137–18148.

Wen H, Hubbard JM, Rakela B, Linhoff MW, Mandel G, Brehm P (2013) Synchronous and asynchronous modes of synaptic transmission utilize different calcium sources. eLife 2:e01206–e01206.

Yu X-M, Askalan R, Keil GJ, Salter MW (1997) NMDA Channel Regulation by Channel-Associated Protein Tyrosine Kinase Src. Science 275:674.

Yu X-M, Salter MW (1998) Gain control of NMDA-receptor currents by intracellular sodium. Nature 396:469–474.

